# Are Current AI Virtual Cell Models Useful for Scientific Discovery?

**DOI:** 10.64898/2026.04.23.719015

**Authors:** Michael Bereket, Jure Leskovec

## Abstract

AI models are increasingly developed to predict the effect of perturbations on gene expression, but current benchmarks fail to reliably measure model performance. Here, we argue that new benchmarks that directly measure the value of model predictions for specific scientific discovery outcomes are needed to address this gap. We present PerturbHD, an evaluation framework for AI-enabled hit discovery, to demonstrate the benefits our proposed approach.

## Main

Understanding the effects of chemical and genetic perturbations on cell states is important for biomedical research, with applications including drug discovery (what molecules revert disease states?) and cell engineering (what genetic modifications induce desired behaviors?). Computational methods that predict perturbation effects, sometimes called AI Virtual Cell models, have been proposed as powerful tools to accelerate scientific discovery by enabling scientists to investigate perturbation effects at a scale infeasible with laboratory experiments [1, 2]. This vision has motivated significant Virtual Cell efforts across academia and industry, with a particular focus on developing models that predict perturbation effects on gene expression [3–11].

However, current benchmarks fail to reliably measure the scientific value of Virtual Cell models. Popular metrics have been shown to score simple constant-prediction baselines as competitive with published deep learning methods, raising concerns regarding both model quality and metric validity [12–14]. While many new metrics have been proposed to address this issue [7, 15–19], no consensus has been reached for effective benchmarking. The ongoing difficulty of defining robust benchmarks was highlighted by the recent Virtual Cell Challenge [2] when participants identified that simply rescaling predictions could improve scores despite increasing prediction error [20]. Without interpretable and trustworthy benchmarks, a critical question remains unanswered: are current AI Virtual Cell models actually useful for scientific discovery?

Here, we argue that current benchmarks are fundamentally limited for characterizing the scientific utility of Virtual Cell models due to a disconnect between the abstract metrics used to measure performance and the capabilities that matter for discovery (Fig. 1a). This manifests as three concrete issues. First, popular metrics are not interpretable in terms of downstream capabilities: for example, does an average correlation of 0.6 mean that a model is a trustworthy tool for experiment design? Second, current metrics fail to provide reliable model rankings. Seemingly intuitive metrics can be misleading: in 3 of 4 popular benchmark screens [21, 22], predicting that all perturbation effect sizes are zero achieves a better mean absolute error (MAE) than effect estimates from experimental replicates (Extended Data Fig. 1, Extended Data Fig. 2). More generally, metrics generate conflicting model rankings and the field lacks principled methods to identify which metrics are meaningful (Extended Data Fig. 1). Finally, standard benchmarks narrow the field by requiring gene-level perturbation effect predictions, excluding promising methods like language models that can directly support downstream discovery needs.

**Figure 1:**
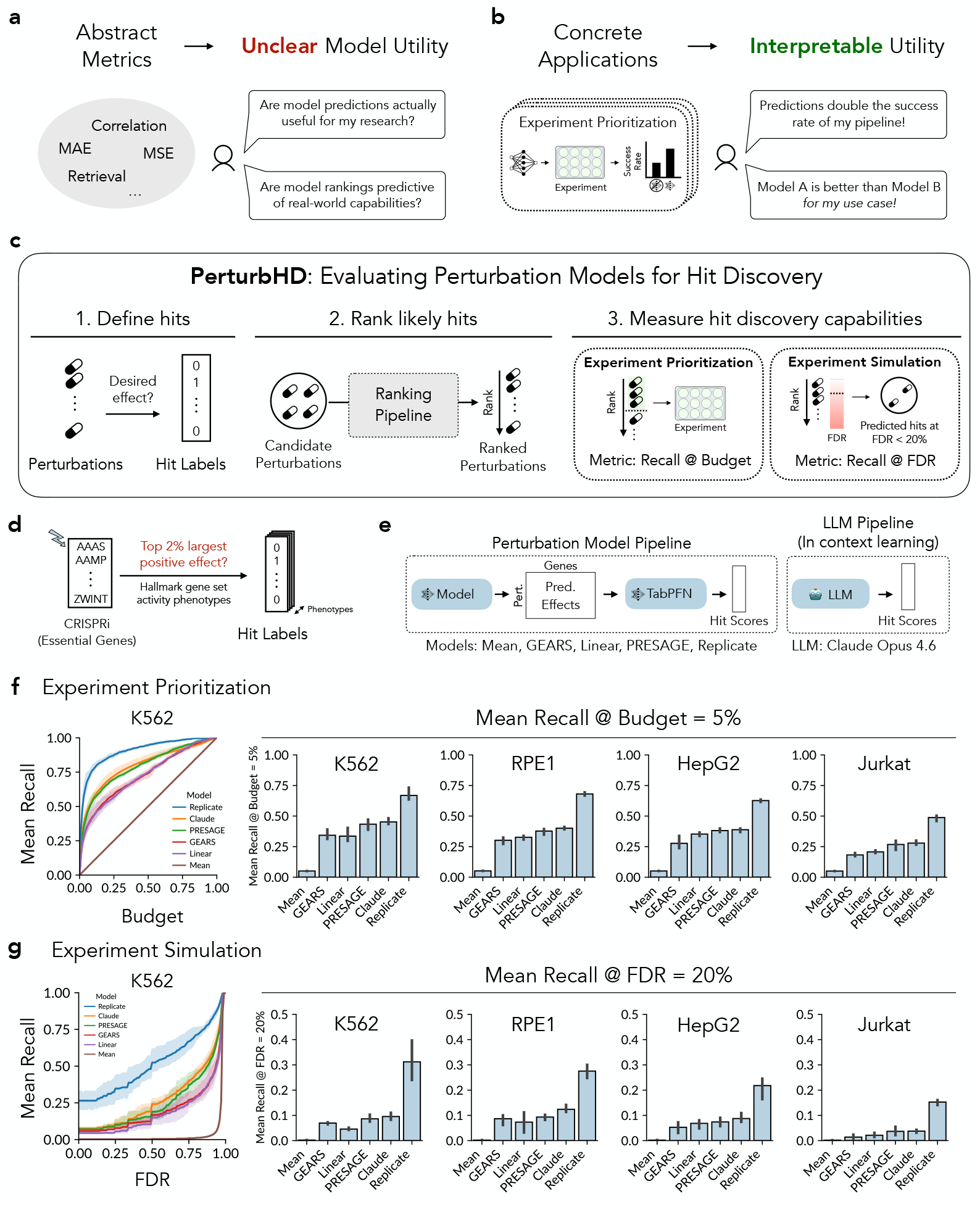
Towards interpretable evaluations of AI Virtual Cell models for scientific discovery. **a)** Popular abstract metrics fail to interpretably measure model performance. **b)** We propose that new metrics that directly measure model utility for specific discovery outcomes can address this gap. **c)** PerturbHD evaluation framework overview. **d–e)** Hit definitions, models, and ranking strategies for experiments. **f–g)** PerturbHD evaluation results for experiment prioritization and simulation in four benchmark datasets. Error bars represent 95% confidence intervals.

We argue that these issues can be solved with new benchmarks that place predictions in concrete scientific workflows and directly assess their impact on scientific discovery outcomes (Fig. 1b). Such evaluations resolve all three limitations discussed above: downstream performance is inherently interpretable, conflicting abstract metrics can be validated against specific discovery capabilities, and the focus on evaluating downstream outcomes rather than intermediate predictions naturally accommodates methods excluded from current benchmarks.

We developed PerturbHD, a framework for evaluating AI-enabled phenotypic hit discovery, to demonstrate our proposed approach. In drug discovery, scientists screen large perturbation libraries to identify *hits*, perturbations with desired phenotypic effects. We analyzed predictions for two hit discovery tasks: experiment prioritization and simulation. In prioritization, predictions are used to rank which experiments to run with the goal of increasing hit discovery efficiency. In simulation, predictions are used to replace experiments by identifying hits directly from predictions while controlling the false discovery rate (FDR). Both tasks can be posed as ranking problems: given a ranked list of likely hits, prioritization considers selecting a fixed budget for testing, while simulation involves selecting a variable number of predicted hits based on the estimated FDR.

PerturbHD evaluates experiment prioritization and simulation in three steps (Fig. 1c). First, a user specifies the desired hit criterion. Next, predictions are used to rank perturbations by hit probability. PerturbHD permits any method that generates a valid ranking, allowing diverse prediction strategies to be compared. Finally, perturbation rankings are evaluated for hit discovery: experiment prioritization is measured as recall at a fixed experiment budget, and experiment simulation is measured as recall at a target empirical FDR.

We applied PerturbHD to analyze model predictions in four large-scale CRISPRi Perturb-seq screens targeting essential genes [21, 22]. Hits were defined as perturbations that strongly upregulate (top 2%) MSigDB Hallmark gene set activity as estimated with AUCell [23, 24] (Fig. 1d). We evaluated three deep learning methods (GEARS [5], linear models with GenePT embeddings [25], and PRESAGE [7]), as well as a constant mean baseline and effect estimates from replicated experiments, using a pipeline that converts gene-level effect predictions to hit rankings using TabPFN [26] (Fig. 1e). As an orthogonal prediction strategy, we evaluated Claude Opus [27], a state-of-the-art LLM, by simply prompting it to rank perturbations given training data as context (Fig. 1e, Extended Data Fig. 4). All models were evaluated using three random splits per dataset, each with approximately 500 unseen test perturbations (~10 hits per phenotype).

We found that PerturbHD provides interpretable insights into the capabilities and limitations of Virtual Cell models for hit discovery. Current models consistently outperformed the mean baseline for experiment prioritization and simulation; however, they were substantially less accurate than experimental replicates (Fig. 1f–g). Interestingly, we found that Claude Opus 4.6 achieved competitive performance with PRESAGE, the best specialized Virtual Cell model, despite no task-specific finetuning or tooling. PerturbHD also goes beyond relative rankings to characterize the overall utility of predictions: while model rankings were similar for prioritization and simulation, the tested Virtual Cell models were far more effective as tools for experiment prioritization (recall 25-45% at a 5% experiment budget) than as experiment simulators (recall 4-12% at 20% FDR). These results highlight key benefits of direct assessments of discovery outcomes: model rankings are interpretable (e.g. an LLM is competitive with PRESAGE *for prioritization in this setting*), a broader class of methods can be compared (the LLM approach would be excluded from standard benchmarks), and assessments of overall utility can support scientific decision making (such as whether to use predictions for prioritization or simulation).

PerturbHD also provides a principled basis to resolve current confusions regarding conflicting metrics. Looking across datasets and models, we found that some existing metrics, like MAE top 100 and Systema correlation [17], were highly predictive of experiment prioritization performance (average Spearman correlation ≥0.95) (Fig. 2, Extended Data Fig. 3). Other metrics like MAE were uncorrelated. While these metric correlations are dependent on the specific set of compared models and do not guarantee universal alignment, careful analyses of the consistency between abstract metrics and desired discovery capabilities can be used to identify metrics that generate biologically meaningful rankings.

**Figure 2:**
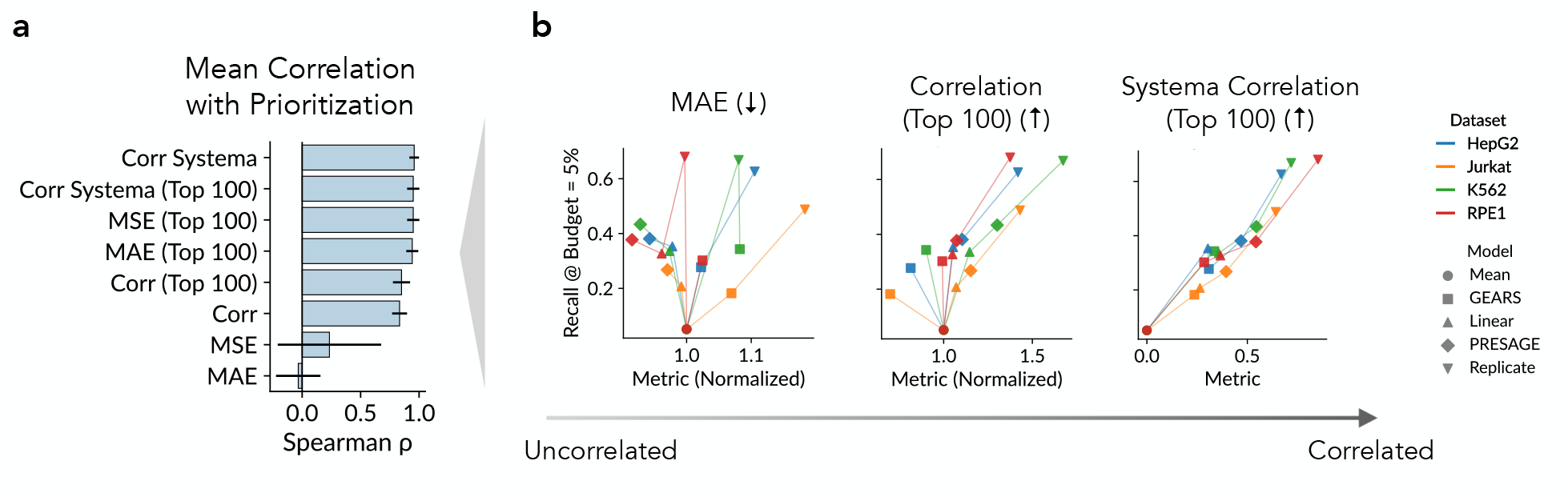
PerturbHD enables principled selection of abstract metrics as proxies for hit discovery performance. **a)** Mean Spearman correlations between model rankings on PerturbHD experiment prioritization and popular abstract metrics. Error bars represent 95% confidence intervals. **b)** Example scatter plots comparing PerturbHD prioritization scores (recall at a 5% budget) and abstract metrics. Each point represents a model-dataset combination, averaged across seeds.

Taken together, these results demonstrate that discovery-oriented evaluations like PerturbHD can address the core limitations of current benchmarks. We note that hit discovery performance will vary across use cases and that our experiments have important differences from standard drug discovery settings: for example, hits are often rarer, perturbation libraries are often larger, and there is often greater distribution shift between training and test perturbations. As a result, PerturbHD should be applied to data and hit definitions tailored to specific applications in order to generate relevant insights.

To conclude, we return to our motivating question: are current AI Virtual Cell models useful for scientific discovery? We argued that existing benchmarks are insufficient to answer this question, and proposed the development of new discovery-oriented metrics to address this gap. In our initial experiments evaluating hit discovery, we observed mixed capabilities: current models showed promise for experiment prioritization, while experiment simulation capabilities were relatively limited. We hope that our analyses encourage biologists and computational researchers to collaborate in creating benchmarks for the wide-range of possible applications of perturbation effect predictions, enabling the field to rigorously characterize the capabilities, limitations, and potential for AI Virtual Cell models for scientific discovery.

## Methods

### Perturb-seq datasets and preprocessing

We uniformly processed four CRISPRi Perturb-seq screens from [21, 22]: K562 essential, RPE1 essential, HepG2 essential, and Jurkat essential. Raw count matrices were processed to remove cells with fewer than 1,000 reads or 500 expressed genes, genes detected in fewer than 5 cells, and perturbations measured in fewer than 20 cells. Counts were normalized to 10,000 counts per cell, log1p transformed, and filtered to a subset of 5,000 highly variable genes using Scanpy (`highly variable genes` with `seurat` method) [28].

We additionally processed the overlapping subset of the K562 genome-wide screen from [21] as a replicate for the K562 essential gene screen. The genome-wide screen was processed identically to the essential gene screens, with the exception of filtering to the same set of genes as the K562 essential screen for consistency. Because only one screen was run in the other cell types, we generated held-out technical replicates by randomly sampling 50% of GEM groups (10x Genomics sequencing batches).

Effect sizes were estimated from Perturb-seq data as the difference in mean log-normalized expression between perturbed cells and control cells receiving non-targeting guides.

### Perturb-seq models

GEARS [5] and PRESAGE [7] were trained with default hyperparameters with the exception of learning rate, which was tuned per dataset. Linear models were fit using Scikit-Learn RidgeCV [29] with GenePT gene embeddings [25] as features and gene-level effect sizes as targets. Effect sizes were standardized per phenotype before fitting, and regularization (*α* ∈ {0.01, 0.1, 1, 10}) was selected by cross-validation. The mean baseline was defined to predict that all perturbations have the same effect, equivalent to the mean effect size vector across training perturbations. All models were fit and evaluated over three random data splits (25% of perturbations held-out as test set).

### Regression and retrieval metrics

We evaluated mean absolute error (MAE), mean squared error (MSE), and Pearson correlation between predicted and estimated effect sizes across all 5,000 gene phenotypes per test perturbation. We also computed “top 100” variants of each metric that restrict comparison to the 100 largest estimated effects per perturbation. Systema correlation variants [17] subtract the average observed perturbation effect vector in the training set from both predicted and observed values before computing correlations, isolating the perturbation-specific component.

### PerturbHD

PerturbHD takes as input a set of test perturbations, binary hit labels, and predicted hit scores, with higher scores reflecting a greater predicted probability that the perturbation is a hit.

Experiment prioritization is evaluated based on recall at experiment budget *b* ∈ [0, 1], where budget *b* corresponds to testing the top ⌈*b*·*n*⌉ of *n* ranked perturbations. Any ties in the perturbation rankings are handled by computing the expected recall over all orderings of the tied group.

Experiment simulation is evaluated based on maximum recall at a target empirical false discovery rate (FDR). Given a ranked list of perturbations, the FDR when selecting the *k* highest-ranked perturbations is FDR(*k*) = (*k* − hits(*k*))*/k*, where hits(*k*) is the number of true hits in the top *k*. For a target FDR *q*, the maximum recall across all cutoffs satisfying FDR(*k*) ≤ *q* is reported. Ties in predicted hit scores are handled by estimating the expected recall over all orderings of tied items using 100 random within-tie permutations.

### PerturbHD experiments

We analyzed perturbation effects on MSigDB Hallmark gene set [23] activity scores computed using AUCell [24] (50 gene sets corresponding to expert-curated biological processes). Gene set activity scores are estimated per cell and average effect sizes are computed as the difference in mean activity between perturbed and control cells. Gene sets are filtered to genes present in the expression matrix.

We converted gene-level predicted effects to hit scores by fitting TabPFN models [26] (a state-of-the-art method for small tabular data) to predict target hit phenotypes from predicted genelevel effects (projected to 100 principal components). Claude Opus 4.6 rankings were generated with a thinking budget of 10,000 tokens by prompting the model to predict hit scores given training data (perturbation, effect size, hit label) as context (Extended Data Fig. 4).

## Data availability

All links for data downloads are available in the provided code repository. K562 Essential, RPE1 Essential, and K562 Genome-Wide screens are downloaded from [21]. Jurkat Essential and HepG2 Essential screens are downloaded from [22].

## Code availability

Code to reproduce all experiments is made available at https://github.com/snap-stanford/perturb-hd.

## Acknowledgements

We thank Yanay Rosen, Moritz Schaefer, Kuan Pang, and Zoe Piran for helpful discussion and feedback.

## Extended Data

**Extended Data Figure 1:**
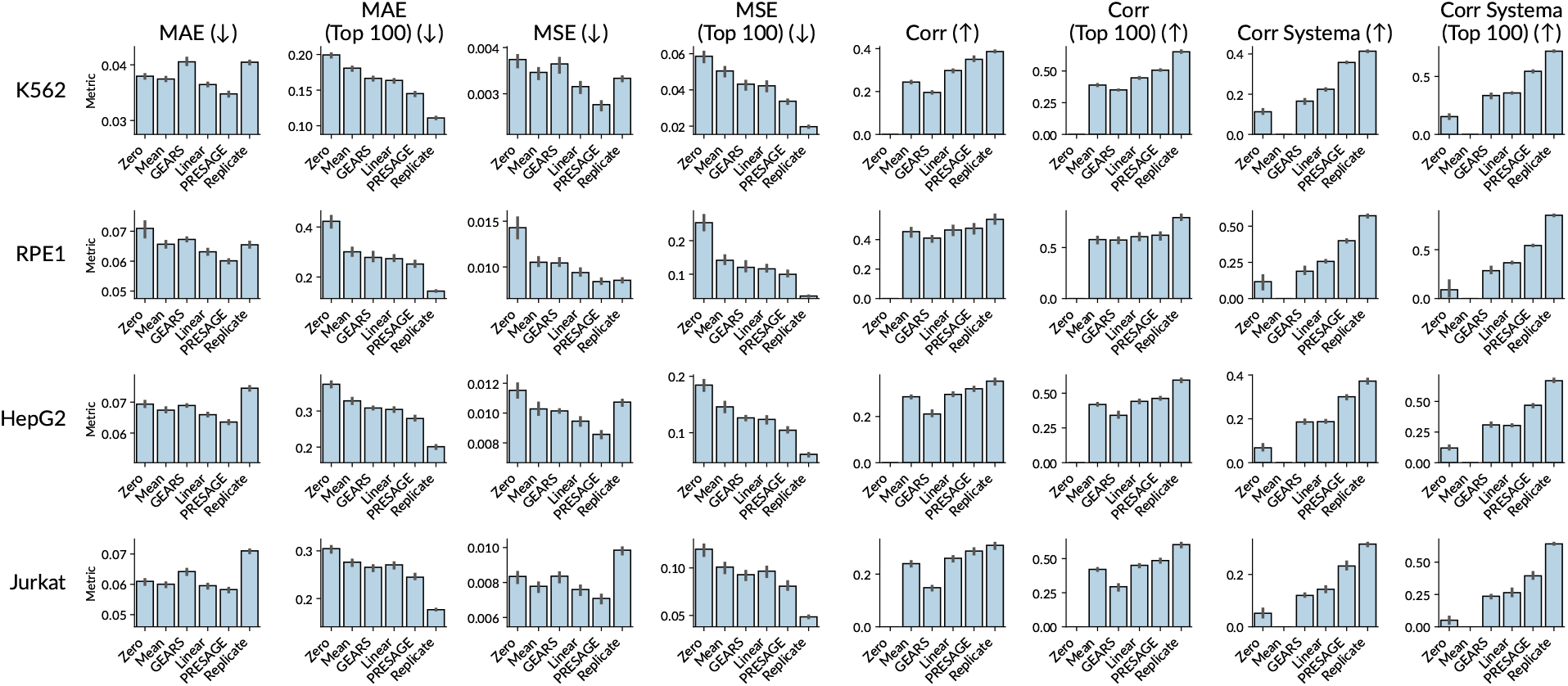
Standard evaluation metrics yield conflicting and difficult to interpret model rankings. Some observations: 1) MAE scores experimental replicates as worse than predicting that all effect sizes are zero in 3 / 4 datasets. 2) MSE scores experimental replicates as worse than or equivalent to PRESAGE, a deep learning model, in all datasets. 3) Metrics disagree on whether GEARS outperforms the mean baseline (correlation and correlation top 100 score the mean baseline as better than GEARS, while MAE top 100, MSE top 100, and both Systema correlation variants score GEARS as better than mean).

**Extended Data Figure 2:**
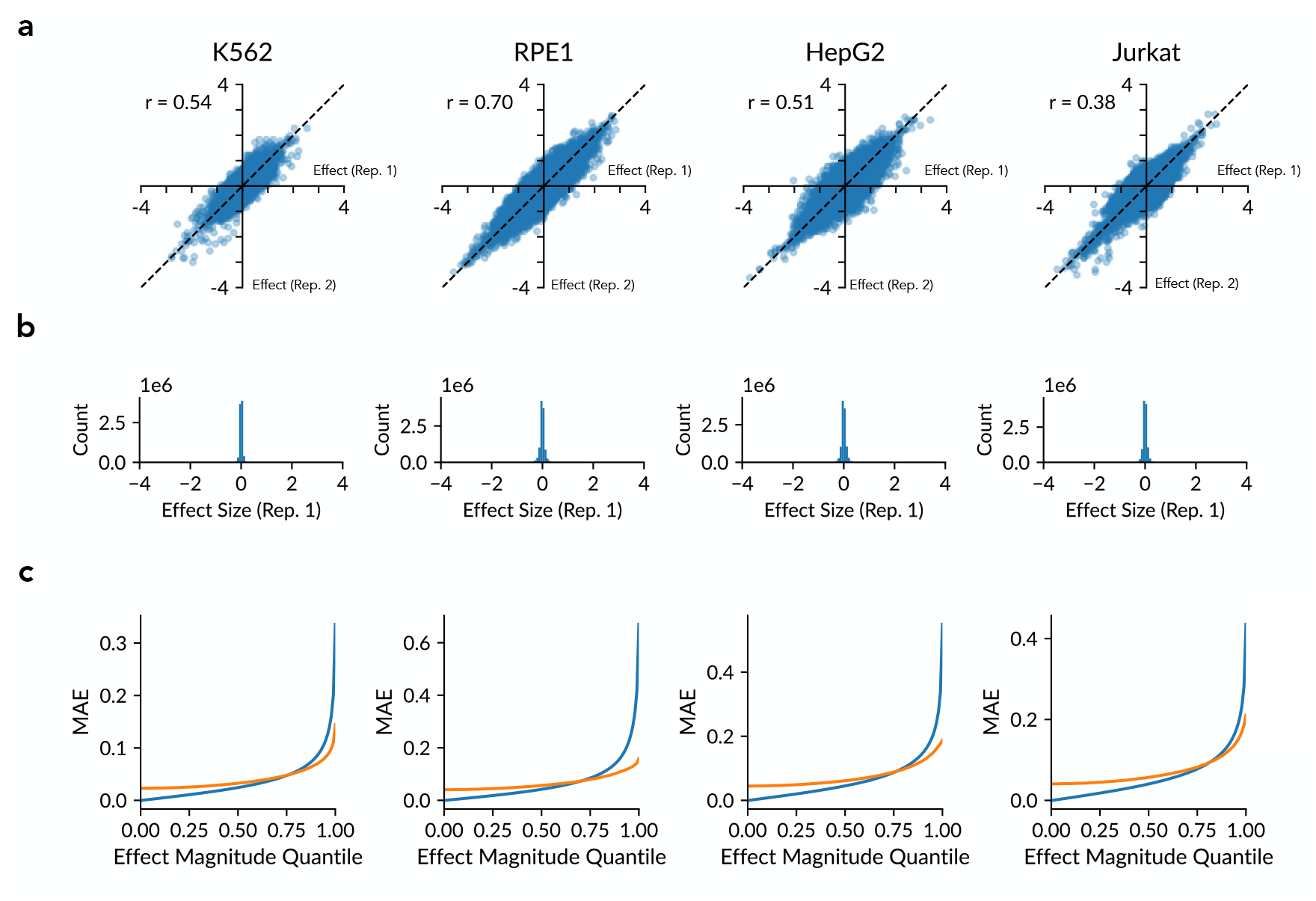
Sparsity and noise in effect size estimates explain the large MAE of replicates. **a)** Effect size estimates are moderately well correlated across replicates. **b)** Effect size estimates are very sparse, with most estimated effects close to zero. **c)** MAE of predicting all effects are zero vs. comparing effect size estimates across replicates, stratified by effect magnitude quantile. For the majority of effects close to zero, estimation error across replicates is larger than the estimated effect magnitudes, resulting in a smaller MAE for all zero predictions in 3 / 4 tested screens.

**Extended Data Figure 3:**
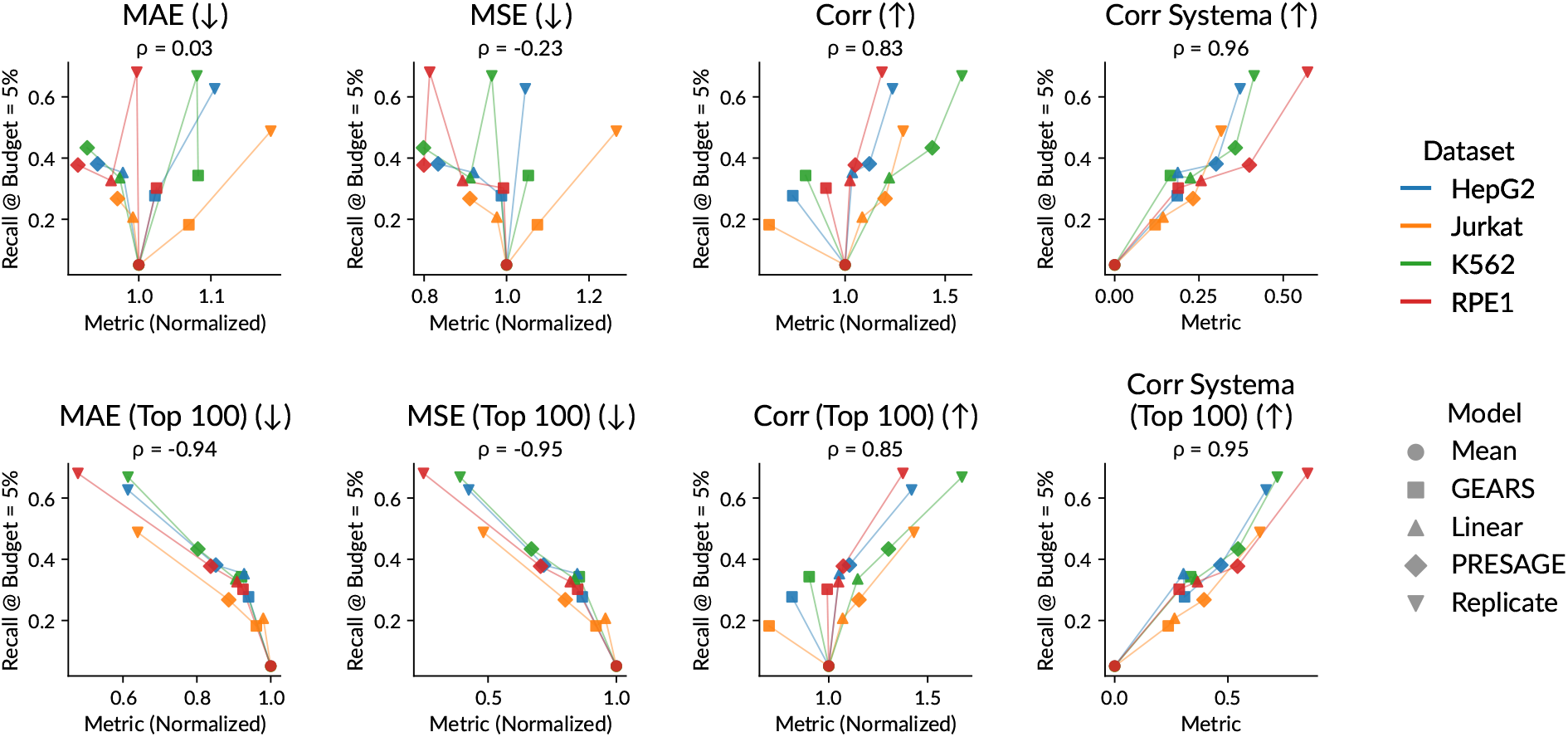
Extended analysis of correlation between PerturbHD and popular abstract metrics. Scatter plots of MAE, MSE, correlation, and Systema correlation scores computed on gene-level effect prediction against prioritization performance (recall @ budget = 5%) across datasets and models. MAE, MSE, and correlation scores are normalized within datasets by dividing by the performance of the mean baseline (Systema correlation is already defined such that train mean is always zero). Average Spearman correlations between each metric and PerturbHD prioritization scores (computed separately for each dataset-seed pair) are indicated above each plot.

**Extended Data Figure 4:**
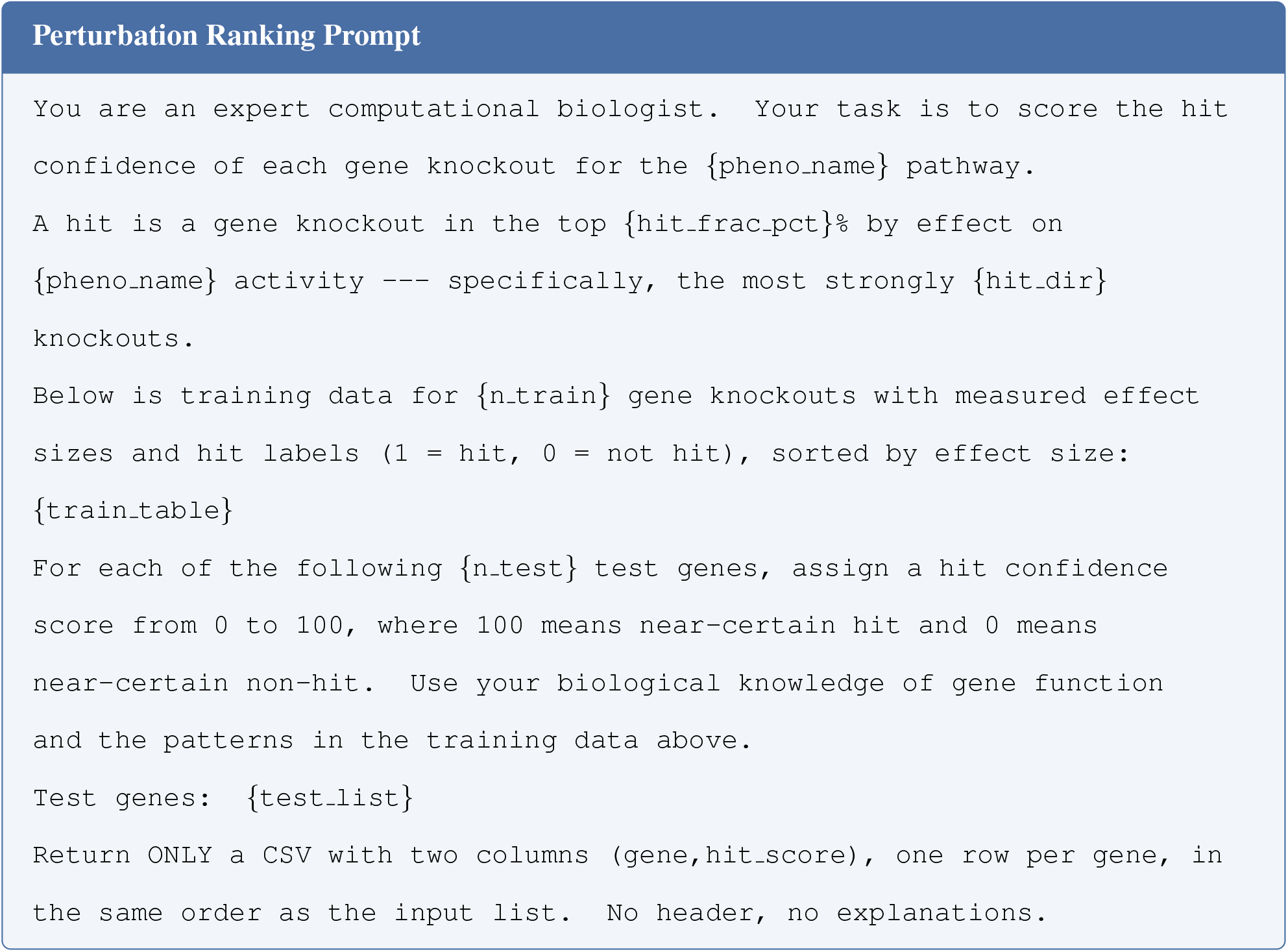
Perturbation ranking prompt. The prompt used to generate LLM-based perturbation rankings for the hit discovery task. Variables surrounded by {} are replaced with values specific to each task. {`train table`} includes perturbation name, effect size on target phenotype, and binary hit labels.

